# Early Reprogramming Intermediates Enable Direct Neuronal Conversion via NGN2

**DOI:** 10.1101/2025.08.04.668497

**Authors:** S Angiolillo, W Qin, A Gesualdo, R Frison, N Elvassore, C Laterza, O Gagliano

## Abstract

The generation of engineered neurons via Neurogenin-2 (NGN2) overexpression, starting from human induced pluripotent stem cells (hiPSCs), is a powerful tool for modeling neurological diseases. However, using stabilized hiPSCs as a starting point significantly increases the time required to obtain a valuable human neuronal model in vitro. Here, we demonstrated that as little as 3 days of transient expression of reprogramming factors in human fibroblasts can unlock the ability of these cells to transdifferentiate into neurons upon overexpression of NGN2. We used single-cell transcriptomic data to dissect the distinct cell identities that emerge during reprogramming. We identified three distinct reprogramming intermediate populations responsive to NGN2-mediated neuronal induction and found that partial reprogramming for only 3 days is sufficient to mediate NGN2 neuronal conversion of human fibroblasts. Moreover, we found that the efficiency in neuronal fate acquisition mediated by NGN2 overexpression is strictly correlated with the stage of reprogramming used as a starting point.

**Highlights:**

- Human fibroblasts undergo reprogramming into pluripotent stem cells, passing through defined intermediate populations.
- Intermediate reprogramming stages can undergo neuronal fate programming by NGN2 overexpression.
- The efficiency of neuronal conversion depends on the intermediate stage employed and increases with reprogramming progression.

## Introduction

The development of reliable human neural in vitro models capable of recapitulating the physiology and pathophysiology of the nervous system is a growing need in the biomedical field. These models should be cost-effective, reproducible, and high-throughput (Nikolakopoulou et al. 2020; Omelchenko, Singh, and Firestein 2020). One of the most efficient and fast methods to obtain human neurons in vitro is through the overexpression of transcription factors in human induced pluripotent stem cells (hiPSCs) or human fibroblasts. Neurogenin-2 (NGN2) is a well-studied proneuronal transcription factor (TF) that is sufficient, alone, to convert hiPSCs into induced neurons (iNs) and to convert human fibroblasts into iNs (Zhang et al. 2013) when used in combination with additional TFs (BRN2, ASCL1, MYTL1, SOX11, SOX4, ISL1, LHX3 and EMX1, according to the neuronal type required) (Hulme et al. 2022) or chromatin modulators (dorsomorphin and forskolin) (Liu et al. 2013). This suggests that NGN2 alone can bind its target regions on hiPSCs but not in differentiated somatic cells, such as fibroblasts, and needs a more accessible chromatin state.

The phenotypic conversion in direct reprogramming protocols, which start from somatic cells (such as fibroblasts) and lead to a different somatic cell type (like neurons), typically requires 3-6 weeks (Hulme et al. 2022), depending on the specific protocol. However, even in the presence of small molecules and miRNAs (Abernathy et al. 2017; Ambasudhan et al. 2011; Victor et al. 2014; Yoo et al. 2011), able to push the differentiation process, the efficiency of direct reprogramming remains extremely low (0.05-5% of the seeded cells) and also requires the cooperation of at least 2-3 transcription factors (Hulme et al. 2022).

On the other hand, the protocols developed for generating iNs starting from hiPSCs, are in general more efficient (50– 60% efficiency, calculated based on the initial number of cells (Zhang et al. 2013)), but extremely time-consuming, if we consider the time required to obtain and stabilize hiPSCs from primary human cells followed by the conversion of hiPSCs to neurons: generating and stabilizing hiPSC lines from human fibroblasts usually requires 4 weeks with an efficiency as low as 0.01-0.1% (Ghaedi and Niklason 2016); generating neurons from hiPSCs usually requires just 2-3 weeks, but can extend to up to 2 months to reach adequate maturation (*i.e*. synapse formation) and functional activity (Togo et al. 2021). Moreover, due to the intrinsic nature of hiPSCs, the same protocols yield different results from different cell lines (Lin et al. 2021), and the presence of any residual pluripotent cells unable to be fully differentiated could alter the meaningfulness of the in vitro model or lead to tumours if used for in vivo applications (Yamanaka 2020).

Recently, to overcome the limitations related to the length of the procedure and the presence of residual pluripotent stem cells, a new approach has been developed. This approach is based on a partial reprogramming followed by neural programming (Capetian et al. 2016; Maza et al. 2015; Meyer et al. 2015). This procedure, known as Pluripotent cell-specific factor-mediated Direct Reprogramming (PDR), uses fully differentiated somatic cells and, by transient expression of pluripotency genes, converts them into pluripotent intermediate cell populations capable of being committed into the cell type of interest (Kim, Ambasudhan, and Ding 2012).

This new method has several advantages, such as the use of a universal pluripotency gene set to generate rejuvenated multipotent progenitor cell populations capable of differentiating into various cell types under specific conditions (Efe et al. 2011; Kim et al. 2011). Furthermore, this strategy significantly reduces the overall timeframe required for cell fate conversion, enabling the transition from fibroblasts to terminally differentiated cell types within approximately 2–3 weeks. Importantly, this method also presents an improved safety profile for in vivo applications; the pluripotent intermediates are not able to proceed toward the acquisition of full pluripotent identity in the absence of exogenously supplied transcription factors, thereby minimizing the risk of tumorigenesis (Yamanaka 2020). Nonetheless, a critical limitation that remains to be addressed for the full realization of this technique’s potential is the currently suboptimal reprogramming efficiency.

In this work, we employed an innovative approach based on the use of partial reprogramming in microfluidics to obtain intermediate cell populations that can be guided to change their identity by transcriptional programming. As we have already shown recently, microfluidics enhances the reprogramming efficiency, leading to precise control over the cellular microenvironment, including both soluble endogenous and exogenous extracellular components, which has a strong positive impact on both the reprogramming (Gagliano et al. 2019; Giulitti et al. 2019; Luni et al. 2016; Panariello et al. 2023) and differentiation processes (Gagliano, Elvassore, and Luni 2016; Luni, Gagliano, and Elvassore 2022).

This partial reprogramming in microfluidics was combined with neuronal programming via direct differentiation into neuronal cells through the induction of the single neural factor NGN2 (Zhang et al. 2013).

We first identified and characterized distinct cellular identities that arise throughout the reprogramming process, analysing single-cell transcriptomic data. This analysis revealed three transcriptionally distinct intermediate populations that exhibit responsiveness to NGN2-driven neuronal induction. Notably, we demonstrated that a brief, 3-day partial reprogramming phase is sufficient to enable NGN2-mediated conversion of human fibroblasts into neurons. Furthermore, our findings indicate that the efficiency of neuronal fate acquisition following NGN2 overexpression is closely dependent on the specific reprogramming stage from which induction is initiated.

This optimised methodology confirms that NGN2 can convert partially reprogrammed somatic cells alone, without additional overexpression of other neuronal TFs.

This strategy can be applied to generate patient-specific in vitro models of iNs in vitro within just 12 days from somatic cells, offering significant potential as a versatile platform for producing human neurons. It holds the promise for advancing regenerative medicine applications, including disease modelling and high-throughput drug screening.

## Results

### Characterisation of reprogramming intermediates

The forced expression of NGN2 alone cannot convert fibroblasts into iNs (Liu et al., 2013), suggesting the need for a more accessible chromatin state to induce neuronal programming. However, there is evidence that a transient reprogramming can unlock the potential of human fibroblasts to become responsive to transcriptional programming (Capetian et al. 2016; Maza et al. 2015; Meyer et al. 2015).

To test whether a partial reprogramming state would allow neuronal programming of human fibroblasts via NGN2 forced expression, we first identified and better characterized the cell populations emerging during a very efficient reprogramming method developed in our lab (Panariello et al. 2023). We demonstrated that a 2-week mRNA-based reprogramming in microfluidics allows a strictly controlled microenvironment, ensuring high efficiency and reproducibility of the reprogramming process (Panariello et al. 2023). This reprogramming method in microfluidics consists of 8 daily transfections of non-modified mRNAs encoding for POU5F1 (OCT3/4), SOX2, KLF4, MYC, LIN28, and NANOG, followed by 7 days of induced pluripotent stem cell (hiPSC) colony stabilisation and expansion in a pluripotency medium (**Fig. 1a, Suppl. Fig. 1a**).

**Figure 1:**
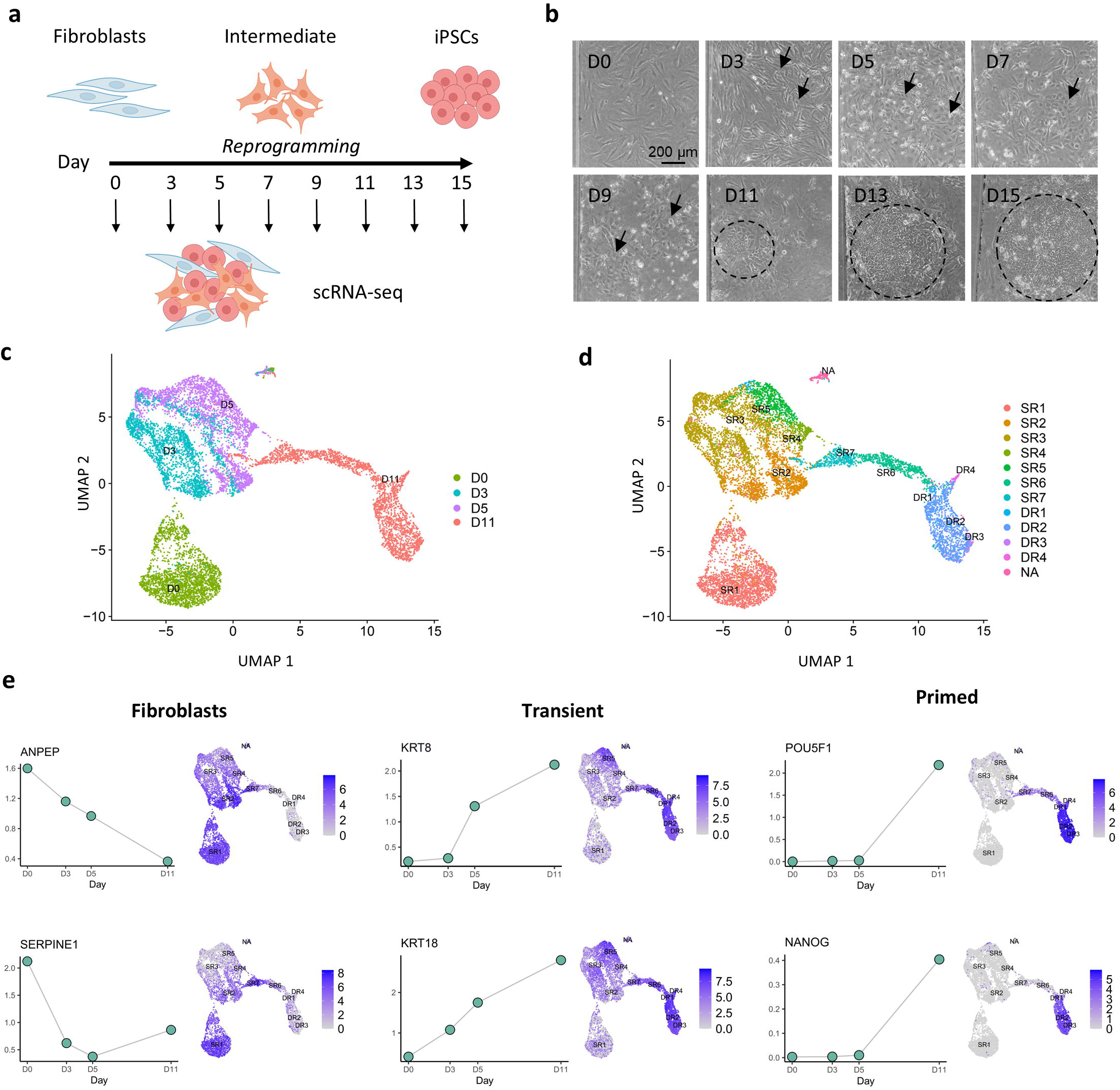
Reprogramming of human fetal fibroblasts in microfluidics generates early intermediate populations undergoing MET remodeling. a) Schematic experimental strategy. b) Morphological conversion along the reprogramming process. Circle: cells undergoing mesenchymal-to-epithelial transition (MET). Scale bar 200 µm. c) UMAP representation of single-cell RNA sequencing data of reprogramming samples collected on days (D) 0, 3, 5, and 11. d) Cluster analysis of single-cell RNA sequencing data. SR: Somatic-related cluster, DR: Developmental-related cluster, NA: non-associated. e) Expression dynamics and UMAP representation showing the expression of fibroblast (left), transient (centre), and primed (right) pluripotency markers in the reprogramming clusters in single-cell RNA sequencing data of D0-3-5-11.

We previously characterised the populations emerging during the process by single-cell RNA sequencing (scRNA-seq) (Panariello et al. 2023) (**Fig. 1a**): cells were assigned to either “Somatic-Related” clusters (SR1-7) corresponding to the states closer to fibroblasts or “Developmental-Related” clusters (DR1-4, with DR4 being a differentiated cluster) corresponding to states that proceeded towards a pluripotent state. Cells transit from an initial state of fibroblasts through a step of intermediate populations that arise between days 3 and 5 (D3-5).

This is consistent with the observation of a quick mesenchymal to epithelial transition (MET) around D3-5 and the appearance of epithelial clusters and hiPSC colony formation by D11 (**Fig. 1b**).

Since the morphological conversion of fibroblasts first appeared at D3, and given the division of secretory and pluripotency trajectories at D5, we better characterised these early intermediate populations. UMAP representation shows the distribution of cells from D0 to D11 (**Fig. 1c**), with some clusters persisting in more time points: SR1 is mostly represented by non-transfected fibroblasts in D1 samples, SR2-3 emerge in D3-5 samples, SR4-5 in D5, while the SR6-7 and the developmental clusters DR1-3 are enriched only in the D11 sample (**Fig. 1d**).

By looking at the evolving expression of fibroblasts, transient (MET and early pluripotency) and late pluripotency markers, we first found by D5 a consistent decrease in fibroblast marker expression, such as ANPEP, SERPINE1, MMP1, and SNAI2 (**Fig. 1e, Suppl. Fig. 1b**), mostly enriched in SR1-2-4 clusters (**Suppl. Fig. 1c**).

Transient markers of MET show an opposite trend, starting to increase around D3-5, such as KRT8 and KRT18, encoding for epithelial cell markers cytokeratin. Similarly, transient early pluripotency markers (PODXL, B3GALT5) are also activated at the D3-5 stage, contributing to cell adhesion and migration. The transmembrane protein PODXL, whose glycosylation by B3GALT5 generates the TRA-1-60 epitope, is a commonly used early marker for undifferentiated human pluripotent stem cells (Lin et al. 2020; Schopperle and DeWolf 2007).

Proper pluripotency markers, such as POU5F1 (OCT3/4), NANOG, KLF5, and SFRP2, show a more restricted expression in DR1-3 clusters on D11, as previously shown (Panariello et al. 2023).

However, it is interesting to note that such transient (PODXL, B3GALT5) and late pluripotency (KLF5, SFRP2) markers are also expressed by SR5-7 (D5-11), other than the expected DR1-3 clusters on the pluripotency trajectory (D11 only) (**Suppl. Fig. 1b**), suggesting that these intermediate populations have a MET phenotype too.

Taken together, these data suggest that the intermediate stage of reprogramming around D3-5 is characterized by subpopulations highly remodelled at the morphological and gene expression levels. They lose their mesenchymal behaviour to acquire a defined intermediate phenotype, expressing transient markers. Just at a later reprogramming stage (D11), the late pluripotency markers emerge, with a defined primed identity.

### The early pluripotent populations are already competent for NGN2 programming

Considering that the D11 sample already shows the activation of some pluripotency markers, we tested whether this late reprogramming stage was sufficient for NGN2 alone to induce neural programming.

We transfected cells with pluripotency factors for 8 days and then cultured them for 3 days in a pluripotency medium; the resulting cell populations should be enriched in cells from clusters SR6-7 and DR1-3. Cells were then transduced at D11 with two lentiviral vectors, pLV-TetO-NGN2-puro and FUW-M2rtTA, expressing NGN2 and Puromycin resistance under the control of Doxicyclin and rtTA, respectively. From the day after the transduction cells were cultured with a neural induction medium containing Doxycycline to activate NGN2 transcription (**Fig. 2a**). As shown in **Fig. 2b-c**, at D11 the populations in the microfluidic chamber were mostly characterised by compact colonies of hiPSCs (positive for OCT3/4, NANOG and SSEA4) surrounded by somatic cells, showing no expression of neuronal markers. After a few days of neural induction (D13-15 from seeding), a morphological change was visible, with previously round cells becoming elongated and losing cell junctions with neighboring cells, ultimately losing their epithelial identity. Then, induced neurons (iNs) with a clear neuronal morphology emerged and matured in complex networks after 9 days of induction (D20 from seeding). At this time point, most of the NGN2^+^ iNs expressed the pan neuronal marker TUJ1/βIIITUB (**Fig. 2d**), except for some compact NANOG^+^ hiPSC colonies and some mesenchymal-like cells.

**Figure 2:**
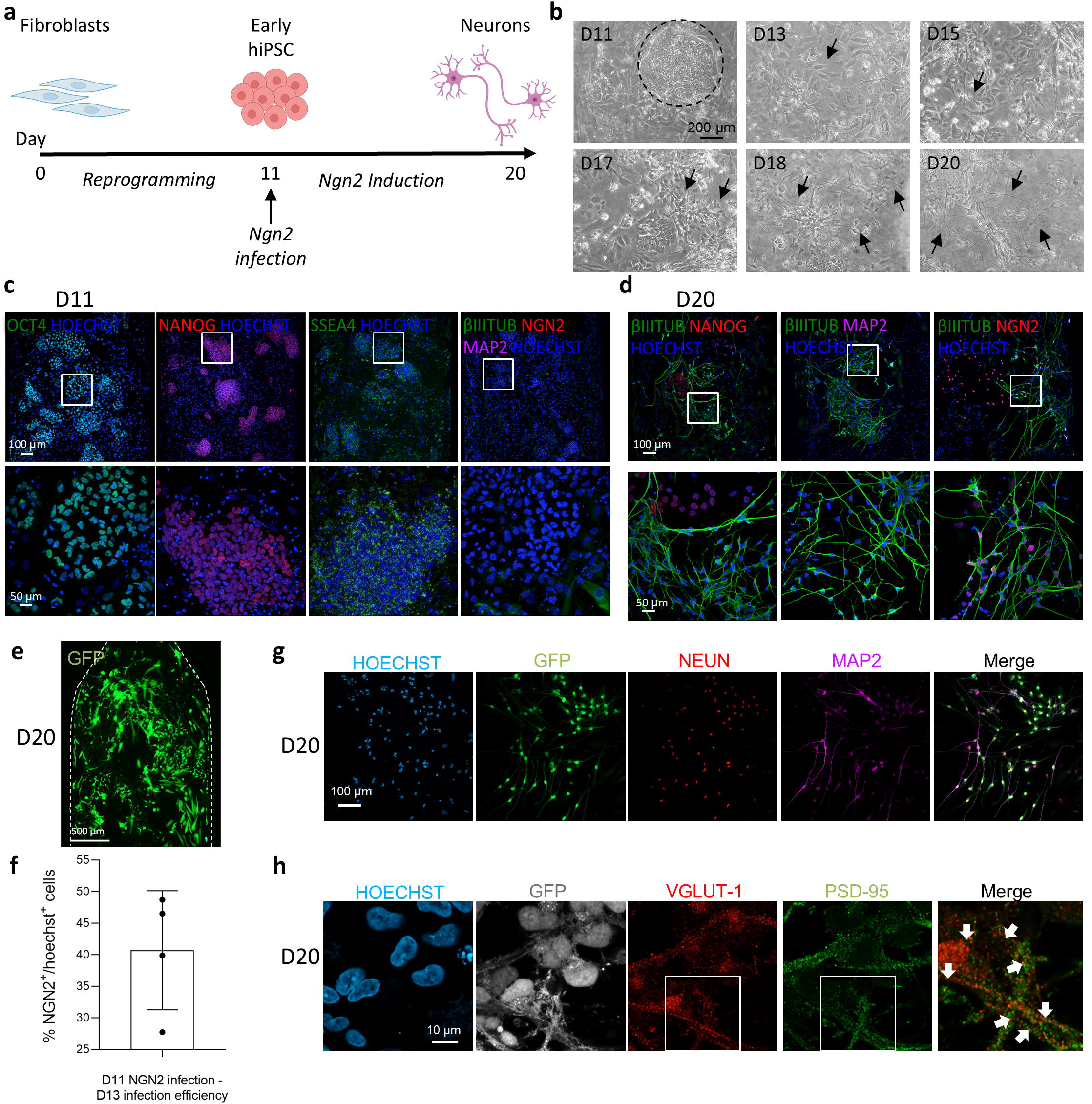
Neuronal conversion via NGN2 induction in early pluripotency populations. a) Schematic experimental strategy. b) Morphological neural conversion after NGN2 induction. Scale bar 200 µm. c) Representative immunofluorescence staining of D11 of reprogramming. Scale bars: 100 μm and 50 μm. d) Representative immunofluorescence staining at D20 of reprogramming-NGN2 programming. Scale bar: 100 µm and 50 µm. e) Representative picture of a microfluidic channel section (dotted line) at D20 of reprogramming-NGN2-eGFP programming. Scale bar: 500 µm. f) Quantification of NGN2^+^ cells over DAPI, at D13 after 2 days of Doxycyclin induction. Expressed as a percentage; median 43.20%, SD: 9.43. g) Representative immunofluorescence staining at D20 of reprogramming-NGN2-eGFP programming of the neural maturation markers NEUN and MAP2. Scale bar: 100 µm. h) Representative immunofluorescence staining at D20 of reprogramming-NGN2-eGFP programming of the synaptic markers VGLUT-1 and PSD-95. Scale bar: 10 µm.

This conversion was not observed in fibroblasts transduced with pLV-TetO-NGN2-puro and FUW-M2rtTA and switched to a neural induction medium containing doxycycline to activate NGN2 transcription for 9 days (**Suppl. Fig. 2a**): although most cells showed overexpression of NGN2, they retained a fibroblast morphology and the expression of mesenchymal markers (VIMENTIN) but not epithelial or neuronal ones (E-CADHERIN and βIIITUB, respectively). This internal control further reinforced the notion that NGN2 alone is not sufficient to transdifferentiate fibroblasts into neurons.

To better track the transduction efficiency, we repeated the same reprogramming-neural programming experiment by using a lentiviral vector (pLV-TetO-NGN2-eGFP-puro) in which the eGFP expression is controlled by Doxicycline administration, as well as for the NGN2 and puromycin resistance genes (**Fig. 2e**).

Two days after NGN2 induction via Doxycycline administration, almost half of the cells expressed eGFP and were positive for NGN2 (median 43.20%, SD: 9.43) (**Fig. 2f, Suppl. Fig. 2b**).

At D20, iNs were distributed in an intricate network with long projections. Moreover, immunofluorescence staining showed that cells were positive for the mature neuronal markers NEUN and MAP2 (**Fig. 2g**) and had co-localized pre-synaptic marker VGLUT-1 and post-synaptic marker PSD-95 (**Fig. 2h**), confirming the presence of glutamatergic synapses and the glutamatergic neuronal fate induced by NGN2 overexpression.

This conversion is not visible when NGN2 overexpression was not induced by Doxycycline supplementation in the neural medium, as shown in the control without Doxycycline by the presence of NANOG^+^ hiPSC colonies and the absence of βIIITUB^+^ neurons (**Suppl. Fig. 2c**).

Our results show that early pluripotent subpopulations emerging at late reprogramming stages are competent to differentiate into neurons by forcing the expression of NGN2 alone in microfluidics. This shows that hiPSCs can be directly programmed toward neural fates without prior stabilization or in vitro expansion.

### Partial reprogramming-neural programming strategy via NGN2 overexpression only

Given that early pluripotent populations are capable of neuronal induction via the overexpression of NGN2, but not fibroblasts, we investigated whether intermediate populations at early reprogramming stages (D3-5) are competent for NGN2 induction.

To answer this question, we transfected cells with the reprogramming factors for just 5 days, instead of the standard 8 (**Fig. 3a**). This partial reprogramming led cells to undergo MET and form small clusters (**Fig. 3b**). These populations were enriched by the SR2-3-4-5 clusters of cells, which we showed to have transient and early pluripotency markers (**Fig. 1e-f**). Then, D5-intermediates were transduced with lentiviral vectors pTet-O-NGN2-puro and FUW-M2rtTA expressing NGN2 and Puromycin resistance under the control of Doxycycline and switched to a neural induction medium supplemented with Doxycycline the following day. After 5 days (D10) of NGN2 induction, we already identified some elongated cells with small nuclei (**Fig. 3b**), positive for βIIITUB and NGN2 (**Fig. 3c**). After 9 days of NGN2 induction (D14), we could observe neuronal cells forming a network with elongated βIIITUB^+^ neurites and some sparse NANOG^+^ pluripotent colonies (**Fig. 3d**), with a neuronal conversion efficiency of 36.4% over the total number of cells (median: 39.13, SD: 17.65), and 64.31% over the initial number of seeded cells (median: 60.74, SD: 21.78) (**Fig. 3e**).

**Figure 3:**
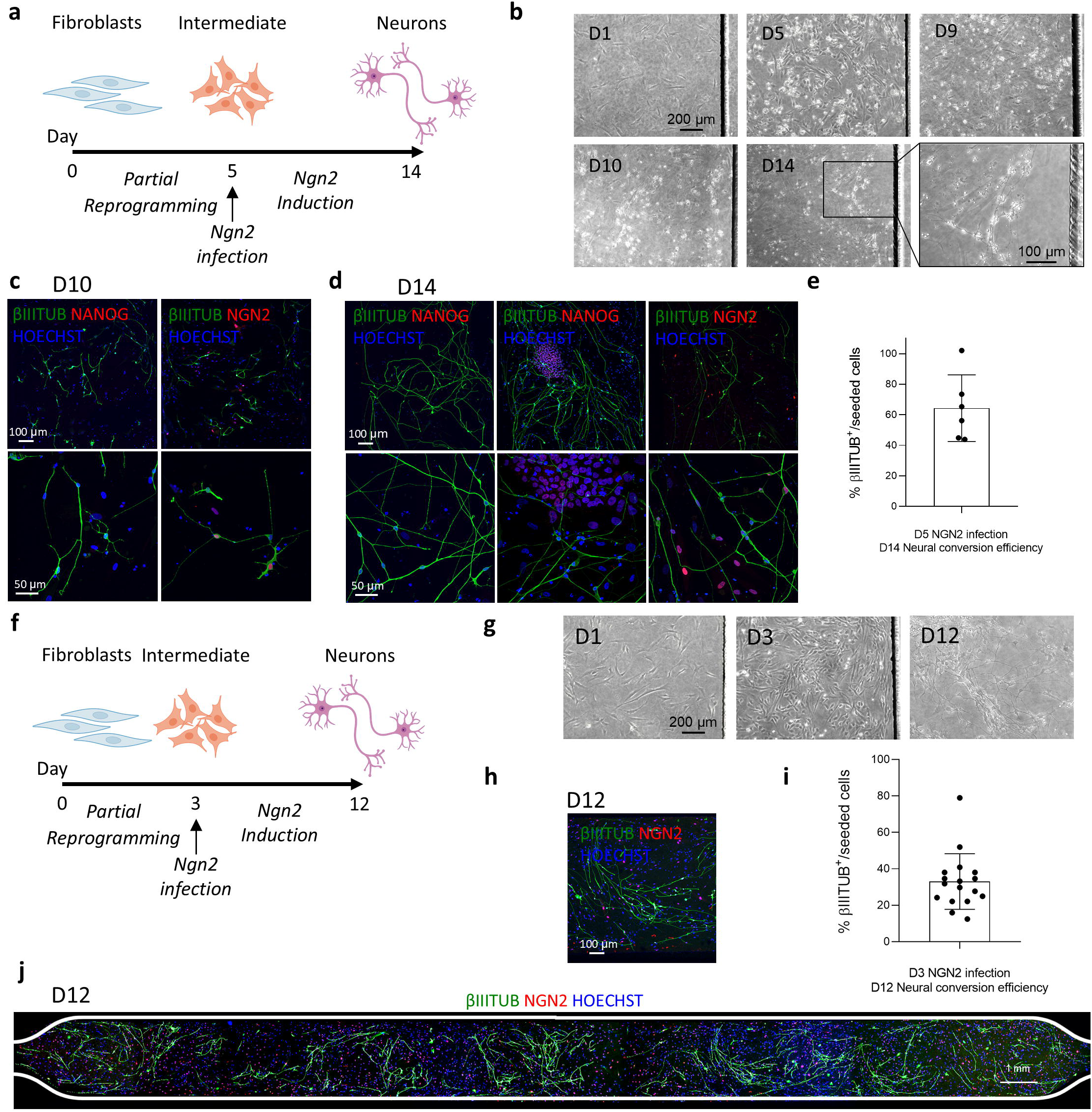
Neuronal conversion based on partial somatic reprogramming and NGN2 induction. a) Schematic experimental strategy of reprogramming, D5 NGN2 infection, and neural programming. b) Morphological neural conversion after partial reprogramming and NGN2 induction. Scale bars: 200 μm and 100 μm. c-d) Immunostaining characterization of D10 and D14 of reprogramming and neuro programming. Scale bars: 100 μm and 50 μm. e) Quantification of neural conversion efficiency at D14 after NGN2 infection at D5 (mean with SD). f) Schematic experimental strategy of reprogramming, D3 NGN2 infection, and neural programming. g) Morphological neural conversion after partial reprogramming and NGN2 induction. Scale bar 200 μm. h) Immunostaining characterization of D12. Scale bar 100 μm. i) Quantification of neural conversion efficiency at D12 after NGN2 infection at D3 (mean with SD). j) Immunostaining characterization of D12 of an entire microfluidic channel. Scale bar 1 mm.

Similar results were obtained by transducing the intermediate reprogramming stage D3 (**Fig. 3f**), mostly enriched in SR2-3 clusters cells and forming round cells in MET, but no visible clusters (**Fig. 3g**). After 9 days of NGN2 induction (D12) we could observe βIIITUB^+^ network of neurons, but no pluripotent colonies (**Fig. 3h**), with a surprisingly high conversion efficiency all along the microfluidic channels of 33.03% over the initial number of seeded cells (median: 31.85, SD: 15.24) (**Fig. 3h-i**). Even though the efficiency of generating iNs from D5-intermediates was almost double that of D3-intermediates, in D3-intermediates, the absence of epithelial colonies was visible at the end of the neural programming, reinforcing the idea that a very early reprogramming intermediate generated by just 3 transfection with pluripotency factors is responsive to NGN2 programming, but not able to undergo complete reprogramming into pluripotent colonies (**Fig. 3h-i-j**).

Our results show that intermediate reprogramming stages, generated after only 3 transfections with reprogramming factors, are compliant with neuronal programming via NGN2 overexpression. This confirms our hypothesis that a short administration of pluripotency factors is sufficient to replace TFs normally cooperating with NGN2 in forcing the neuronal fate acquisition in somatic cells.

## Discussion

The present study demonstrates that transient exposure to reprogramming factors for as little as three days is sufficient to enable human fibroblasts to acquire responsiveness to NGN2-mediated neuronal programming. Our data indicate that subpopulations emerging during intermediate reprogramming stages undergo a profound transcriptional and phenotypic remodeling, marked by MET, downregulation of fibroblast-specific genes, and activation of early pluripotency-associated markers. These findings suggest that such intermediate states, obtained at day 3 and 5, create a permissive chromatin landscape that enables efficient direct conversion into neurons upon NGN2 induction, without the need for full stabilization into hiPSCs.

Previous reports have shown that direct conversion of fibroblasts into induced neurons (iNs) by transcription factor overexpression is limited by low efficiency and requires the combination of multiple proneural genes or chromatin-modulating small molecules (Liu et al. 2013; Pfisterer et al. 2016). In contrast, the overexpression of NGN2 alone is sufficient to induce neuronal fate from fully stabilized hiPSCs, but fails when applied directly to somatic fibroblasts (Liu et al. 2013). Our findings support the hypothesis that the permissiveness to NGN2 action depends on the chromatin state and transcriptional plasticity of the starting population (Soufi et al. 2015). By implementing a microfluidic-based protocol for transient reprogramming, we were able to reproducibly generate intermediate populations with features of MET and early pluripotency markers that allow efficient NGN2-mediated neuronal induction.

Single-cell transcriptomic analysis revealed that reprogramming intermediates generated by day 3 to day 5 display the greatest heterogeneity and harbour transient expression of epithelial and early pluripotency genes such as PODXL and B3GALT5, which are typically involved in cell adhesion and glycosylation processes associated with stem cell identity (Lin et al. 2020; Schopperle and DeWolf 2007). Notably, we observed that even minimal exposure to reprogramming factors (three daily mRNA transfections) is sufficient to activate these pathways and enable the acquisition of neuronal fate upon NGN2 induction. The lack of stable hiPSC colony formation in D3-derived populations, despite their competence for neuronal programming, further supports the notion that full reprogramming to pluripotency is not a prerequisite for NGN2-mediated neuronal induction. Moreover, a deeper investigation into the identity and characteristics of these D3-derived populations may reveal a transient intermediate state that is capable of homogeneously transdifferentiating into neurons without generating fully pluripotent cells. Interestingly, the conversion efficiency varied with the reprogramming stage, being highest in D11 freshly reprogrammed iPSCs and decreasing from D5 to D3 intermediates, suggesting that there exists an optimal temporal window during which cells are most permissive to neuronal fate induction. This aligns with observations in other reprogramming systems where transient activation of pluripotency circuits enhances cellular plasticity without fully erasing somatic identity (Meyer et al. 2015; Mitchell et al. 2014).

The application of microfluidic technology was instrumental in enabling neuronal fate induction in D3 intermediate populations. This platform played a critical role in standardizing the reprogramming microenvironment, thereby improving the reproducibility and efficiency of phenotypic conversion.

By controlling medium flow and ensuring a stable biochemical cell niche, microfluidics supports efficient mRNA delivery and minimizes variability across experiments, as we also previously reported (Luni et al. 2016; Panariello et al. 2023). From a research perspective, this approach presents multiple advantages. First, it reduces the time and cost associated with the generation and stabilization of hiPSC lines. Second, it mitigates the tumorigenic risks associated with residual pluripotent cells in downstream applications (Yamanaka 2020). Third, the ability to generate patient-specific iNs within 12 days opens the way for personalized disease modeling and high-throughput screening in neurodegenerative and neurodevelopmental disorders (Lin et al. 2021; Zhang et al. 2020). Nevertheless, further studies will be required to assess the functional maturity, electrophysiological properties, and long-term stability of iNs derived from early reprogramming intermediates.

In conclusion, our work uncovers a narrow temporal window of cellular plasticity during partial reprogramming, in which somatic cells gain competency for NGN2-driven neuronal fate acquisition.

This “reprogramming-to-programming” strategy represents a rapid and efficient method to generate human neurons, providing new insights into the interplay between transient pluripotency circuits and lineage-specific programming. Future efforts will focus on optimizing the epigenetic landscape of intermediate populations and extending this approach to the generation of specific neuronal subtypes relevant for disease modeling and drug screening studies.

**Suppl Figure 1: Reprogramming of human fetal fibroblasts in microfluidics generates early intermediate populations undergoing MET remodeling**. a) Representative immunofluorescence staining of a microfluidic channel on D15 at the end of the reprogramming process, showing the co-expression of the pluripotency markers TRA1-60 and NANONG on hiPSC colonies. b) Expression dynamics and UMAP representation showing the expression of fibroblast (left), transient (centre) and primed (right) pluripotency markers in the reprogramming clusters in single-cell RNA sequencing data of D0-3-5-11. c) UMAP representation showing the expression of fibroblast (first column), transient (second column) and primed pluripotency (third column) markers in the reprogramming clusters.

**Suppl Figure 2: Neuronal conversion requires both NGN2 induction and responsive cellular phenotype**. a) (Left) schematic experimental strategy of fibroblast infection with lentiviruses expressing pLV-TetO-NGN2-puro and FUW-M2rtTA but no neural conversion. (Right) Representative immunofluorescence staining at D9 of NGN2 induction. Scale bars: 100 µm and 50 µm. b) (Left) schematic experimental strategy of infection with lentiviruses expressing pLV-TetO-NGN2-puro and FUW-M2rtTA at D11 of reprogramming, and 2-day NGN2 induction. (Right) Representative immunofluorescence staining at D13. Scale bar 100 µm. c) (Left) schematic experimental strategy of infection with lentiviruses expressing pLV-TetO-NGN2-puro and FUW-M2rtTA at D11 of reprogramming, and no NGN2 induction. (Right) Representative immunofluorescence staining at D20. Scale bar: 100 µm and 50 µm.

## Author Contributions

O.G., C.L., and N.E. designed the study. S.A. and W.Q. optimized and performed the reprogramming and neural programming experiments, with the help of R.F.. C.L. and S.A. produced the lentiviral vectors. S.A. performed immunofluorescence stainings, with the help of R.F., and image quantifications. A.G. analyzed the single-cell RNA sequencing data. W.Q. and R.F. fabricated the microfluidic devices. O.G., C.L., and S.A. critically discussed the data and wrote the manuscript. O.G. and C.L. supervised the project.

## Acknowledgements

This research was supported by the University of Padova under the 2019 STARS Grants program (iNeurons to O.G.) and 2021 STARS Grants program (EnCOr to C.L.).

## Methods

### Microfluidic device

In this work, we used a microfluidic device, fabricated by soft lithography technique and replica moulding, previously published by our group (Gagliano et al. 2019). Polydimethylsiloxane (PDMS) with a 10:1 base/curing agent ratio (Dow Corning) was coupled to a borosilicate glass slide (Menzel–Gläser) through plasma treatment of surfaces.

The standard height of each channel is 200 µm, and the area is 27 mm^2^. Each channel holds 10 µL of medium when considering the inlet, the outlet, and the culture channel itself. The device is sterilized by autoclaving before use. During experiments, the microfluidic chips are placed in a dish, surrounded by a water bath with PBS -/- to reduce medium evaporation.

### Reprogramming in microfluidics

The reprogramming protocol was performed from human foreskin BJ fibroblasts (ATCC) as previously described (Gagliano et al. 2019; Panariello et al. 2023). On day 0 (D0), microfluidic chambers are coated with 25 µg/mL VTN-N (Vitronectin, ThermoFisher A14700) for 1 h at room temperature, and fibroblasts are seeded at 60 cells/mm2 in DMEM (Gibco, 41965) + 10% Fetal Bovine Serum (FBS; Gibco, 10270106).

On D1 morning, the medium is switched to the reprogramming medium (Nutristem hPSC XF Medium, Biological Industries 06-5100-01-1A, supplemented with 50 ng/mL FGF-2, Peprotech 100-18B-1000). On D1 afternoon, transfections are performed using StemRNA-NM reprogramming kit (Stemgent, 00-0076) and StemMACS mRNA transfection kit (Miltenyi, 130-104-463) in the reprogramming medium.

These steps are repeated for 8 days in case of the full reprogramming protocol, and on D9 the medium is replaced with hiPSC medium (StemMAC iPS-Brew XF, Miltenyi Biotec, 130-107-086). In case of the partial reprogramming protocols, transfections have been reduced to 5 or 3.

The whole reprogramming process is performed at a medium change frequency of 12 h in hypoxic conditions (5% O2, 5% CO2) at 37°C.

### Lentiviral vector production

All lentiviral vectors were handled in a class II biosafety laboratory. All plasmids were purchased on Addgene: pLV-TetO-NGN2-puro (Addgene, 52047), pLV-TetO-NGN2-eGFP-puro (Addgene, 79823), FUW-M2rtTA (Addgene, 20342), pRSV-REV(Addgene, #12253), pMDLg/pRRE (Addgene, #12251), and pMD2.G (Addgene, #12259). 293-T cells (Invitrogen) were seeded in T175 flasks (Sarstedt), previously coated with Poly-Ornithine (10 µg/ml in water) overnight at room temperature, in DMEM with Glutamax (Gibco 10565018) + 10% v/v FBS (Gibco A5256701) + 1x Non-Essential Amino Acids (NEAA, Gibco 11140-035) + 1x Sodium Pyruvate (Gibco, 11360-070). When they reached 75-90% confluency, plasmids containing the transgenes of interest were co-transfected with pMDL/RRE (Addgene, 12251), pRSVREV (Addgene, 12253), pMD2G (VsVG, Addgene, 12259), and PEI (1 μg/ml, Polysciences, 23966-1) into the 293-T cells to initiate viral particle production. The medium was changed 14-16h after transfection, supernatants were harvested at 48 h after transfection and ultracentrifuged at 20 000 x g for 2h at 4°C to concentrate the lentiviral particles. Pellets were resuspended in PBS overnight, aliquoted and stored at -80°C.

### Neural programming in microfluidics

At the different stages of the reprogramming (D3-5-11 of reprogramming), the cells have been co-transduced with NGN2-expressing lentiviral vectors (pLV-TetO-NGN2-puro or pLV-TetO-NGN2-eGFP-puro) and FUW-M2rtTA lentiviral vectors according to the protocol published by Zhang et al. 2013 with some modifications. The day after, neuronal phenotype was induced by switching to a neural induction medium supplemented with Doxycycline (2.5 μg/mL final concentration, Clontech) to induce the TetO gene and puromycin to select the population infected, for 9 days. Neural induction is composed of Advanced DMEM/F12 (Gibco, 12634-010): Neurobasal medium (1:1; Gibco, 21103-049), 0.25× N2 (Gibco, 17502048), 0.5× B27 (Gibco, 17504044), 1× Non-essential Amino Acids (Gibco, 11140-035), 2 mM l-glutamine (Gibco, 25030024), and 50 µM β-mercaptoethanol (Gibco, 31350-010). For the last 4 days, the medium is further supplemented with Brain-derived neurotrophic factor (BDNF) (20 ng/ml, Peprotech, 450-02) and Neurotrophin-3 (NT3) (20 ng/ml, Peprotech, 450-03).

### Immunofluorescent staining

Microfluidic chambers were fixed in 4% paraformaldehyde (Sigma, P6148) for 10 minutes at room temperature, and then washed three times with PBS -/-. Samples were blocked with 5% Donkey serum (Merck, S30) in 0.1% Triton X-100 (PBST) at room temperature for 1 h. Primary antibodies were diluted in the blocking solution and incubated overnight at 4°C. Then, samples were washed three times with 0.1% PBST, incubated with secondary antibodies in the blocking solution for 1 h at room temperature, washed three times with 0.1% PBST, and then PBS -/-.

The following antibodies were used for immunofluorescence: rabbit anti-NANOG (Cell Signaling, 4903, 1:200), mouse anti TRA1-60 (Milli-pore, MAB4360, 1:100), mouse anti-OCT3/4 (Santa Cruz, sc-5279, 1:200), mouse anti-SSEA4 (Santa Cruz, sc-21704, 1:250), mouse anti-TUJ1/βIIITUB (Biolegend, 801213, 1:5000), rabbit anti-NGN2 (Cell Signalling, 13144S, 1:100), chicken anti-MAP2 (Abcam, Ab5392, 1:8000), chicken anti-GFP (Millipore, 06-896, 1:300), rabbit anti-NEUN (Abcam, AB104225, 1:500), rabbit anti-V-GLUT1 (SYSY, 135303, 1:100), mouse anti-PSD-95 (Abcam, AB2723, 1:200), rabbit anti-VIMENTIN (Sigma, SAB1305445-40TST, 1:50), mouse anti-E-CADHERIN (BD, 610181, 1:50).

Alexa488, Alexa594, and Alexa647 conjugated rabbit and mouse secondary antibodies were used (Life Technologies, 1:500), or Alexa488/Alexa647 conjugated chicken secondary antibodies (Jackson Immunoresearch, 1:500). The nuclei were stained with Hoechst (Life Technologies, 33342).

Images were acquired on a confocal TCS SP5 microscope (Leica) at 40x magnification and on a fluorescence microscope DM6B (Leica) at 5 and 10x magnification.

The efficiency of lentiviral infection was calculated after 2 days, by expressing the ratio of NGN2^+^ cells over the total number of Hoechst^+^ cells.

The neural conversion efficiency of human fibroblasts into neurons was calculated by counting the TUJ1/βIIITUB^+^ and Hoechst^+^ cells per visual field. The area of each visual field was then measured, and we used this estimated density of iNs and the total amount of cells to determine the total number of neurons present in the entire channel.

We then divided this number by the number of cells plated before starting the reprogramming to obtain the percentage of the starting population of the cells that adopted neuron-like characteristics.

### scRNA-seq analysis

The scRNA-seq dataset used in this paper was published by Panariello et al. 2023, and the raw data can be found in the GEO database under accession code GSE221739. Only four time points of reprogramming of the published dataset were retained for the purpose of this paper: D0, D3, D5, and D11. The analysis of the dataset was performed using the R Seurat package (v. 5.1.0) in RStudio under R version 4.4.0. Values of gene expression are normalised to CPM and transformed to the log2 scale using a pseudocount of 1 to standardize the expression counts. Principal Component Analysis (PCA) on the scaled data was computed, and non-linear dimension reduction using uniform manifold projection (UMAP) was performed using 14 principal components. Cluster identification and annotation were retained from the original paper. The expression of selected marker genes associated with fibroblasts, primed iPSCs, and transient pluripotency was shown across the samples using the function FeaturePlot implemented in the Seurat package. The variation over time of the expression of the same marker genes was also shown after pseudo bulk aggregation of the cells in each time point using the AggregateExpression function in the Seurat package.

## References

Abernathy, D. G., Kim, W. K., McCoy, M. J., Lake, A. M., Ouwenga, R., Lee, S. W., Xing, X., Li, D., Lee, H. J., Heuckeroth, R. O., Dougherty, J. D., Wang, T., & Yoo, A. S. (2017). MicroRNAs Induce a Permissive Chromatin Environment that Enables Neuronal Subtype-Specific Reprogramming of Adult Human Fibroblasts. Cell Stem Cell, 21(3), 332-348.e9. 10.1016/j.stem.2017.08.002

Ambasudhan, R., Talantova, M., Coleman, R., Yuan, X., Zhu, S., Lipton, S. A., & Ding, S. (2011). Direct Reprogramming of Adult Human Fibroblasts to Functional Neurons under Defined Conditions. Cell Stem Cell, 9(2), 113–118. 10.1016/j.stem.2011.07.002

Capetian, P., Azmitia, L., Pauly, M. G., Krajka, V., Stengel, F., Bernhardi, E.-M., Klett, M., Meier, B., Seibler, P., Stanslowsky, N., Moser, A., Knopp, A., Gillessen-Kaesbach, G., Nikkhah, G., Wegner, F., Döbrössy, M., & Klein, C. (2016). Plasmid-Based Generation of Induced Neural Stem Cells from Adult Human Fibroblasts. Frontiers in Cellular Neuroscience, 10. 10.3389/fncel.2016.00245

Efe, J. A., Hilcove, S., Kim, J., Zhou, H., Ouyang, K., Wang, G., Chen, J., & Ding, S. (2011). Conversion of mouse fibroblasts into cardiomyocytes using a direct reprogramming strategy. Nature Cell Biology, 13(3), 215–222. 10.1038/ncb2164

Gagliano, O., Elvassore, N., & Luni, C. (2016). Microfluidic technology enhances the potential of human pluripotent stem cells. Biochemical and Biophysical Research Communications, 473(3), 683–687. 10.1016/j.bbrc.2015.12.058

Gagliano, O., Luni, C., Qin, W., Bertin, E., Torchio, E., Galvanin, S., Urciuolo, A., & Elvassore, N. (2019). Microfluidic reprogramming to pluripotency of human somatic cells. Nature Protocols, 14(3), 722–737. 10.1038/s41596-018-0108-4

Ghaedi, M., & Niklason, L. E. (2016). Human Pluripotent Stem Cells (iPSC) Generation, Culture, and Differentiation to Lung Progenitor Cells (pp. 55–92). 10.1007/7651_2016_11

Giulitti, S., Pellegrini, M., Zorzan, I., Martini, P., Gagliano, O., Mutarelli, M., Ziller, M. J., Cacchiarelli, D., Romualdi, C., Elvassore, N., & Martello, G. (2019). Direct generation of human naive induced pluripotent stem cells from somatic cells in microfluidics. Nature Cell Biology, 21(2), 275–286. 10.1038/s41556-018-0254-5

Hulme, A. J., Maksour, S., St-Clair Glover, M., Miellet, S., & Dottori, M. (2022). Making neurons, made easy: The use of Neurogenin-2 in neuronal differentiation. Stem Cell Reports, 17(1), 14–34. 10.1016/j.stemcr.2021.11.015

Kim, J., Ambasudhan, R., & Ding, S. (2012). Direct lineage reprogramming to neural cells. Curr Opin Neurobiol, 22(5), 778–784. 10.1016/j.physbeh.2017.03.040

Kim, Janghwan, Efe, J. A., Zhu, S., Talantova, M., Yuan, X., Wang, S., Lipton, S. A., Zhang, K., & Ding, S. (2011). Direct reprogramming of mouse fibroblasts to neural progenitors Supporting Information. Proceedings of the National Academy of Sciences of the United States of America, 108(19).

Lin, H.-C., He, Z., Ebert, S., Schörnig, M., Santel, M., Nikolova, M. T., Weigert, A., Hevers, W., Kasri, N. N., Taverna, E., Camp, J. G., & Treutlein, B. (2021). NGN2 induces diverse neuron types from human pluripotency. Stem Cell Reports, 16(9), 2118–2127. 10.1016/j.stemcr.2021.07.006

Lin, R.-J., Kuo, M.-W., Yang, B.-C., Tsai, H.-H., Chen, K., Huang, J.-R., Lee, Y.-S., Yu, A. L., & Yu, J. (2020). B3GALT5 knockout alters glycosphingolipid profile and facilitates transition to human naïve pluripotency. Proceedings of the National Academy of Sciences, 117(44), 27435–27444. 10.1073/pnas.2003155117

Liu, M.-L., Zang, T., Zou, Y., Chang, J. C., Gibson, J. R., Huber, K. M., & Zhang, C.-L. (2013). Small molecules enable neurogenin 2 to efficiently convert human fibroblasts into cholinergic neurons. Nature Communications, 4(1), 2183. 10.1038/ncomms3183

Luni, C., Gagliano, O., & Elvassore, N. (2022). Derivation and Differentiation of Human Pluripotent Stem Cells in Microfluidic Devices. Annual Review of Biomedical Engineering, 24, 231–248. 10.1146/annurev-bioeng-092021-042744

Luni, C., Giulitti, S., Serena, E., Ferrari, L., Zambon, A., Gagliano, O., Giobbe, G. G., Michielin, F., Knöbel, S., Bosio, A., & Elvassore, N. (2016). High-efficiency cellular reprogramming with microfluidics. Nature Methods, 13(5), 446–452. 10.1038/nmeth.3832

Maza, I., Caspi, I., Zviran, A., Chomsky, E., Rais, Y., Viukov, S., Geula, S., Buenrostro, J. D., Weinberger, L., Krupalnik, V., Hanna, S., Zerbib, M., Dutton, J. R., Greenleaf, W. J., Massarwa, R., Novershtern, N., & Hanna, J. H. (2015). Transient acquisition of pluripotency during somatic cell transdifferentiation with iPSC reprogramming factors. Nature Biotechnology, 33(7), 769–774. 10.1038/nbt.3270

Meyer, S., Wörsdörfer, P., Günther, K., Thier, M., & Edenhofer, F. (2015). Derivation of Adult Human Fibroblasts and their Direct Conversion into Expandable Neural Progenitor Cells. Journal of Visualized Experiments, 101. 10.3791/52831

Mitchell, R. R., Szabo, E., Benoit, Y. D., Case, D. T., Mechael, R., Alamilla, J., Lee, J. H., Fiebig-Comyn, A., Gillespie, D. C., & Bhatia, M. (2014). Activation of Neural Cell Fate Programs Toward Direct Conversion of Adult Human Fibroblasts into Tri-Potent Neural Progenitors Using OCT-4. Stem Cells and Development, 23(16), 1937–1946. 10.1089/scd.2014.0023

Nikolakopoulou, P., Rauti, R., Voulgaris, D., Shlomy, I., Maoz, B. M., & Herland, A. (2020). Recent progress in translational engineered in vitro models of the central nervous system. Brain, 143(11), 3181–3213. 10.1093/brain/awaa268

Omelchenko, A., Singh, N. K., & Firestein, B. L. (2020). Current advances in in vitro models of central nervous system trauma. Current Opinion in Biomedical Engineering, 14, 34–41. 10.1016/j.cobme.2020.05.002

Panariello, F., Gagliano, O., Luni, C., Grimaldi, A., Angiolillo, S., Qin, W., Manfredi, A., Annunziata, P., Slovin, S., Vaccaro, L., Riccardo, S., Bouche, V., Dionisi, M., Salvi, M., Martewicz, S., Hu, M., Cui, M., Stuart, H., Laterza, C., … Elvassore, N. (2023). Cellular population dynamics shape the route to human pluripotency. Nature Communications, 14(1), 2829. 10.1038/s41467-023-37270-w

Pfisterer, U., Ek, F., Lang, S., Soneji, S., Olsson, R., & Parmar, M. (2016). Small molecules increase direct neural conversion of human fibroblasts. Scientific Reports, 6(1), 38290. 10.1038/srep38290

Schopperle, W. M., & DeWolf, W. C. (2007). The TRA-1-60 and TRA-1-81 Human Pluripotent Stem Cell Markers Are Expressed on Podocalyxin in Embryonal Carcinoma. Stem Cells, 25(3), 723–730. 10.1634/stemcells.2005-0597

Soufi, A., Garcia, M. F., Jaroszewicz, A., Osman, N., Pellegrini, M., & Zaret, K. S. (2015). Pioneer Transcription Factors Target Partial DNA Motifs on Nucleosomes to Initiate Reprogramming. Cell, 161(3), 555–568. 10.1016/j.cell.2015.03.017

Togo, K., Fukusumi, H., Shofuda, T., Ohnishi, H., Yamazaki, H., Hayashi, M. K., Kawasaki, N., Takei, N., Nakazawa, T., Saito, Y., Baba, K., Hashimoto, H., Sekino, Y., Shirao, T., Mochizuki, H., & Kanemura, Y. (2021). Postsynaptic structure formation of human iPS cell-derived neurons takes longer than presynaptic formation during neural differentiation in vitro. Molecular Brain, 14(1), 149. 10.1186/s13041-021-00851-1

Victor, M. B., Richner, M., Hermanstyne, T. O., Ransdell, J. L., Sobieski, C., Deng, P.-Y., Klyachko, V. A., Nerbonne, J. M., & Yoo, A. S. (2014). Generation of Human Striatal Neurons by MicroRNA-Dependent Direct Conversion of Fibroblasts. Neuron, 84(2), 311–323. 10.1016/j.neuron.2014.10.016

Yamanaka, S. (2020). Pluripotent Stem Cell-Based Cell Therapy—Promise and Challenges. Cell Stem Cell, 27(4), 523– 531. 10.1016/j.stem.2020.09.014

Yoo, A. S., Sun, A. X., Li, L., Shcheglovitov, A., Portmann, T., Li, Y., Lee-Messer, C., Dolmetsch, R. E., Tsien, R. W., & Crabtree, G. R. (2011). MicroRNA-mediated conversion of human fibroblasts to neurons. Nature, 476(7359), 228–231. 10.1038/nature10323

Zhang, Ying, Xie, X., Hu, J., Afreen, K. S., Zhang, C.-L., Zhuge, Q., & Yang, J. (2020). Prospects of Directly Reprogrammed Adult Human Neurons for Neurodegenerative Disease Modeling and Drug Discovery: iN vs. iPSCs Models. Frontiers in Neuroscience, 14, 546484. 10.3389/fnins.2020.546484

Zhang, Yingsha, Pak, C., Han, Y., Ahlenius, H., Zhang, Z., Chanda, S., Marro, S., Patzke, C., Acuna, C., Covy, J., Xu, W., Yang, N., Danko, T., Chen, L., Wernig, M., & Südhof, T. C. (2013). Rapid Single-Step Induction of Functional Neurons from Human Pluripotent Stem Cells. Neuron, 78(5), 785–798. 10.1016/j.neuron.2013.05.029

