## Supplementary figures and images for "Early Reprogramming Intermediates Enable Direct Neuronal Conversion via NGN2"

### Supplementary Figure 1

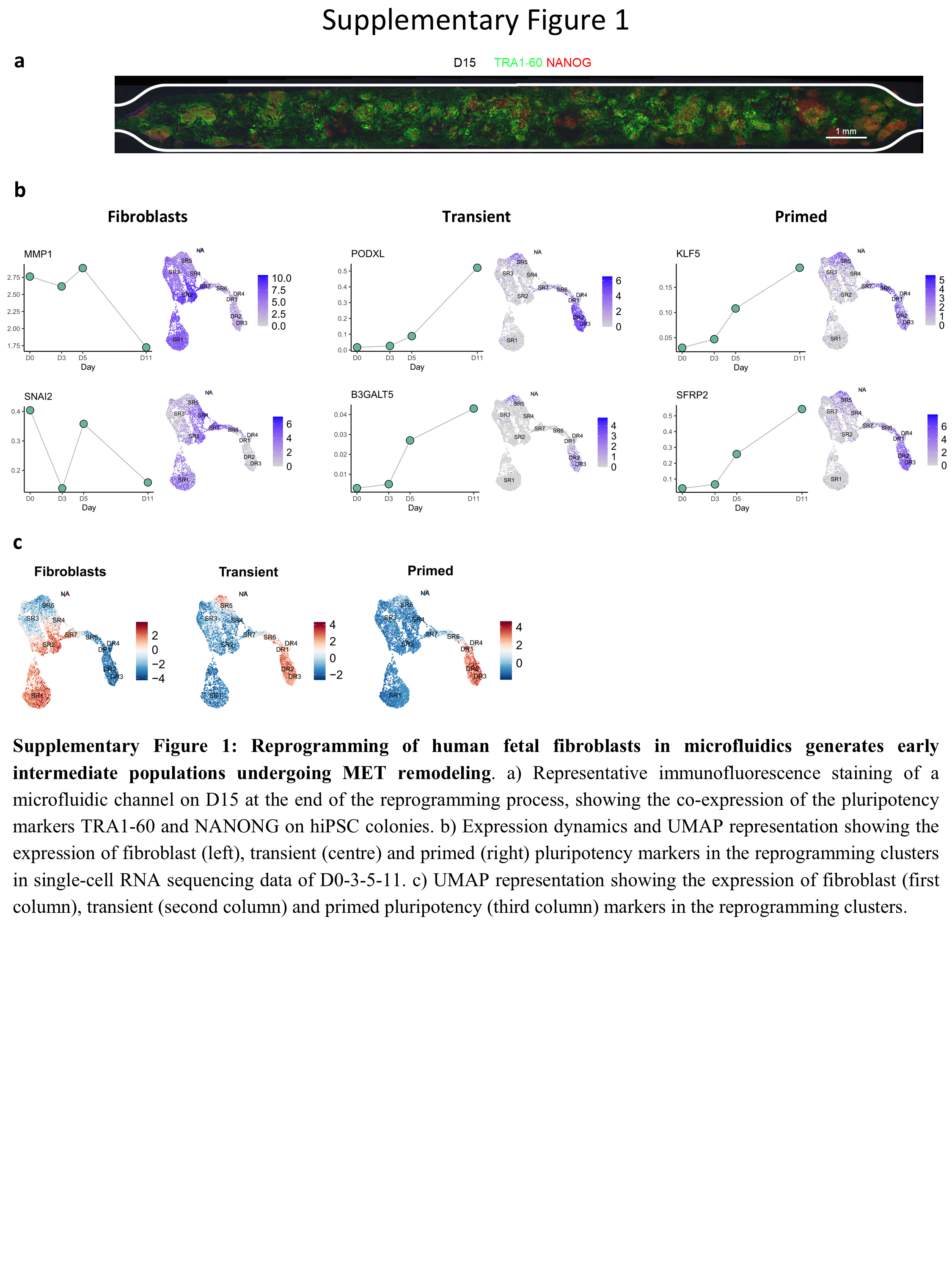

### Supplementary Figure 2

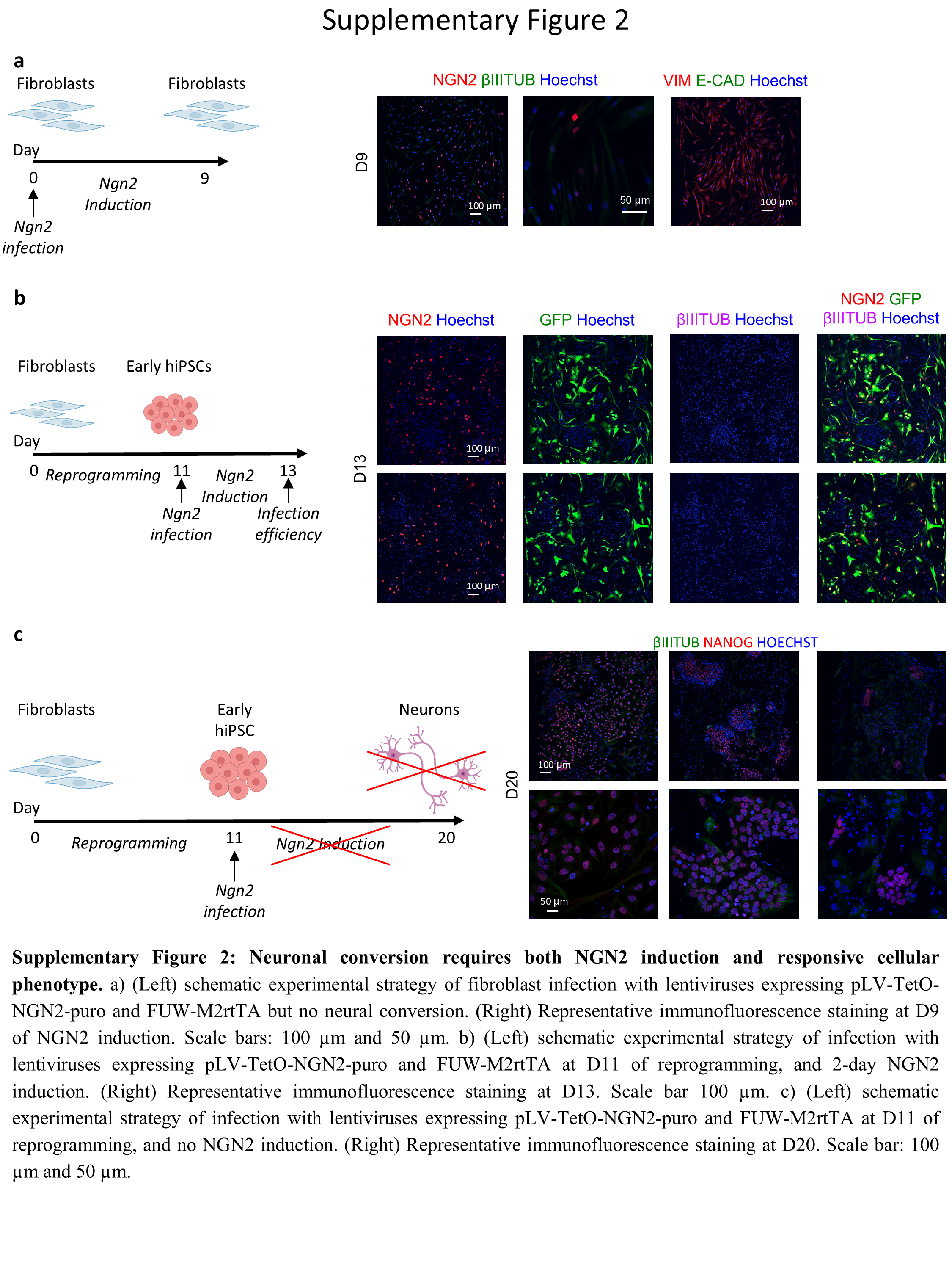
